# Outer hair cell electro-mechanical power conversion

**DOI:** 10.1101/2021.07.29.454372

**Authors:** Richard D. Rabbitt

**Affiliations:** Biomedical Engineering, Otolaryngology, and Neuroscience Program, University of Utah, 36 S. Wasatch Drive, SMBB 3100, Salt Lake City, UT 84112

**Keywords:** electromotility, prestin, cochlea, piezoelectricity, electro-mechanics, biological motor, imaginary capacitance

## Abstract

Outer hair cells are the cellular motors in the mammalian inner ear responsible for sensitive high-frequency hearing. Motor function requires expression of the protein prestin in the OHC lateral membrane and mechano-electrical transduction in the apical hair bundle. In the present report, electrical power consumption and mechanical power output are determined using previously published voltage clamp data from isolated OHCs and membrane patches. Results reveal that power output by the prestin-motor complex peaks at a best frequency much higher than implied by the low-pass character of nonlinear capacitance, and much higher than the whole-cell resistive-capacitive corner frequency. High frequency power output is enabled by a −90° shift in the phase of prestin-dependent electrical charge displacement, manifested electrically as emergence of imaginary-valued nonlinear capacitance.

## Introduction

The sensitivity and frequency bandwidth of mammalian hearing relies on active amplification of sound-induced vibrations in the cochlea by electro-motile outer hair cells (OHCs), which change length in response to changes in membrane potential [1]. Voltage-driven electro-motility is imparted by expression of the transmembrane protein prestin [2, 3] through a mechanism that couples electrical charge displacement in the prestin motor complex to changes in cell length. Electro-mechanical coupling works in reciprocal directions [4], consistent with piezoelectric materials where electrical and mechanical terms are directly coupled in the Gibbs free energy [5]. Piezoelectric equations are thermodynamically self-consistent, and commonly used to describe electro-mechanical coupling in OHCs [6–10]. The structure of prestin [11–13] suggests piezoelectricity on the whole-cell level arises from a large number of prestin dimers undergoing conformational changes on the nanoscale, with each conformational change involving an electrical charge displacement and a change in membrane area occupied by the dimer. In this view, the transmembrane electrical field generates Coulomb forces on charged residues and associated ions which combine to induce a subsequent conformational change in the configuration of the molecular complex.

There are conflicting reports addressing the speed of the process. Optical coherence tomography measurements in the living cochlea support cycle-by-cycle amplification at frequencies up to at least 20 kHz in mouse [14], with amplification occurring primarily near the traveling wave peak in gerbil and mouse [15, 16]. Analysis of power output by OHCs based on vibrations of the cochlear partion [17] provide additional evidence supporting the cycle-by-cycle amplification hypothesis. Other evidence including low-pass character of OHC displacement in echolocating bats [18] and 2^nd^-order distortion products in gerbil [19] challenge the cycle-by-cycle amplification hypothesis. Low-pass characteristics are also present in nonlinear capacitance (NLC) recorded in isolated membrane patches [20–22], and in voltage-driven displacement of isolated OHCs measured using the μ-chamber approach [23, 24]. Precisely how the prestin motor complex amplifies vibrations in the cochlea given these low-pass features remains unknow.

Computational models of cochlear mechanics that treat OHCs as cycle-by-cycle as piezoelectric actuators [8] replicate experimental data reasonably well over a broad bandwidth, including changes in vibrational patterns from apex to base [25]. Theoretical analysis of isolated OHCs driving a mechanical load further support the hypothesis that OHCs inject power into cochlear viscous load cycle-by-cycle over the entire auditory frequency bandwidth [9, 10]. These cycle-by-cycle computational and theoretical considerations are consistent with cochlear amplification in whales and bats at frequencies greater than 100kHz, but upon first inspection seem at odds with the low-pass characteristics of the prestin motor complex.

The present report re-examines data from isolated membrane patches and whole cells under voltage clamp conditions to determine if the data support the hypothesis that power output and electro-mechanical power conversion are low pass. Results reject the null hypothesis and demonstrate that electro-mechanical power conversion is ultrafast, and generates mechanical power output up to the highest frequency tested to date. Peak power output occurs at frequencies much higher than might be expected on the basis of NLC or voltage-driven displacement. Results further support the hypothesis that power output in isolated membrane patches and OHCs is bandpass and dependent on mechanical load, similar that measured in the cochlea [16] and predicted previously on theoretical grounds for isolated cells [9, 10].

## Methods

### Electrical Power Consumed Under Ideal Voltage Clamp

To find the electrical power consumed by the prestin motor complex for small sinusoidal signals under voltage-clamp conditions, the transmembrane voltage, current and electrical charge displacement are written as 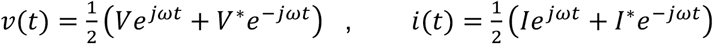 and 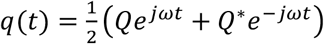, respectively. Lower case denotes the time domain, upper case denotes the frequency domain, the * denotes the complex conjugate and ω is frequency in *rad* – *s*^-1^. The charge *Q* is equal to the capacitance *C* times voltage *V*, where *C* is the sum of the linear capacitance *C^L^* and the prestin-dependent nonlinear capacitance *C^N^*. Multiplying the electrode current *i*(*t*) times voltage *ν*(*t*) and taking the time average gives the electrical power delivered by the patch pipette: 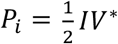. Multiplying the membrane electrical displacement current 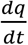 by voltage *ν*(*t*) and taking the time average gives the electrical power *P_E_* consumed by the prestin motor complex:

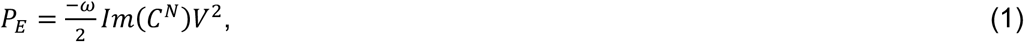

which, by the second law of thermodynamics, must be converted to heat plus mechanical power output by the motor. Notably, *P_E_* is completely independent of the real part of the nonlinear capacitance, demonstrating that *Re*(*C^N^*) not related to mechanical power output by prestin in membrane patches or in OHCs [10].

### Electrical Power Consumed Under General Conditions

To determine the electrical power consumed from experimental data under more general conditions, the OHC is treated as a single electrical compartment, where the voltage and current are related by Kirchhoff’s current law: 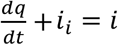. The input current is *i*. and the total ionic channel conduction current is *i_i_*. Under isothermal conditions, the charge displacement depends on voltage *ν* and mechanical strain *s*. From the chain rule of calculus 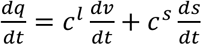, where 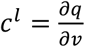 is the capacitance voltage susceptibility (conventional “linear” electrical capacitance) and 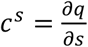 is the capacitance strain susceptibility arising from the prestin motor complex [10, 26]. The voltage *ν* is the transmembrane voltage, which can arise from both intracellular and extracellular voltage modulation in the cochlear organ of Corti. The whole cell displacement *x* is related to the strain by *x* = *ℓs*, where *ℓ* is cell length. For small perturbations from the resting state (*ν_o_*, *i_o_*, *x_o_*), the Fourier transform of Kirchoff’s current balance for small perturbations in the frequency domain gives:

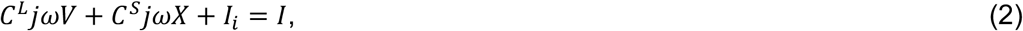

where 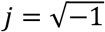. The parameter *C^S^* is proportional to the capacitance strain susceptibility and quantifies the relationship between the electrical displacement current in the prestin motor complex and the whole-cell axial displacement. *I_i_* describes the linearized ionic currents in the frequency domain, while *I* is the input current *via* the MET channels (or electrode).

Electrical power is determined by multiplying Eq. 2 by the complex conjugate of voltage (*V*\*) and taking half of the real part, which is equivalent to the time-domain method used to derive Eq. 1. The right-hand side gives the electrical power input *P_E_ via* the MET channels or patch electrode:

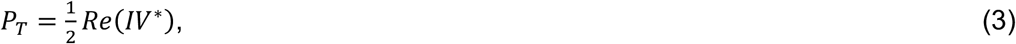

which is the *input electrical power* available from cycle-by-cycle modulation of the voltage and the MET receptor current (or electrode current). *V*\* is the transmembrane voltage modulation determined from potentials on both sides of the membrane, a distinction that is important in the organ of Corti *vs*. recordings from isolated cells in the dish. The left-hand side gives the electrical power lost to ion-channels:

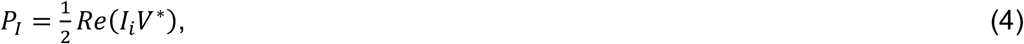

and the electrical power consumed by the prestin-motor complex:

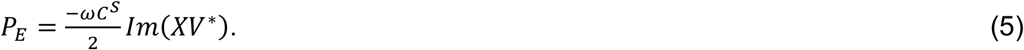

Notably, from Eq. 5, the electrical power consumed by the motor complex is zero if the voltage modulation is in phase with the displacement, and maximum if the voltage is in phase with velocity. This fact means the power consumed depends on the mechanical load acting on the prestin motor complex, because the load alters the displacement *X* even if the voltage *V* is constant. The parameter *C^S^* varies with reference state (*ν_o_*, *i_o_*, *x_0_*) and configuration of prestin, but like passive electrical capacitance *C^L^*, is constant for small sinusoidal perturbations in voltage, current and displacement about the reference state. *C^S^* can be determined experimentally in the frequency domain after blocking ion channels. The direct approach is to measure *V*, *I* and *X* and apply Eq. 2 to determine *C^S^*.

Although simple, the form of Eq. 5 is difficult to use experimentally because it requires measuring *V*, *I* and *X* at the same time. Measuring *P_E_* is simplified under voltage clamp (Eq. 1). From thermodynamics, the two must be identical (*P_E_* from Eq. 1 must be equal to *P_E_* from Eq. 5). Equivalency is demonstrated by writing the OHC displacement as a function of voltage and force: *X* = *X*(*V*, *F*) and expanding in a Taylor series to find the total capacitive current arising from prestin *jωC^S^X* = *jω*(*C^V^V* + *C^F^F*), where the capacitance voltage susceptibility is 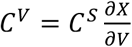 and the capacitance force susceptibility is 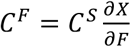. After blocking ion channels, *C^V^* can be measured in voltage clamp under constant force, and *C^F^* can be measured in force clamp under constant voltage. While *C^S^* is a constant independent of load, both *C^V^* and *C^F^* depend on the mechanical strain in the prestin-motor complex and therefore depend on how the cell is loaded. Under ideal voltage clamp, a command voltage *V* causes the cell to displace against the load imposed by the cell itself and the environment. When working against a load, the force is a frequency-dependent function of voltage (e.g. *F* = *TV*, where *T* is a transfer function). Combining terms, the total capacitive current is *jωC^N^V*, where *C^N^* = *C^V^* + *TC^F^*, and the power consumed by the prestin-motor is found to be exactly the same as Eq. 1. Application of Eq. 1 simply requires experimental measurement of *C^N^* and *V*, and is a valid approximation regardless of the biophysical origin(s) of NLC. Eq. 1 clearly has experimental advantages, but Eq. 5 offers more insight into power output and consumption by the prestin motor complex because the capacitance displacement susceptibility *C^S^* is independent of load, while *C^N^* is variable and depends on how the membrane is loaded under conditions of the experiment.

### Mechanical Power Output

The mechanical power output by the prestin-motor complex is time-averaged force times velocity, and can be written in the frequency domain as:

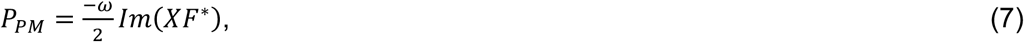

where *X* is the axial cell displacement and *F* is the axial somatic force. Eq. 7 can be used to estimate power output irrespective of the biophysical origins of force, but it is informative to demonstrate how power output is related to electrical power consumed based on thermodynamics of electromechanics which, to first order, is piezoelectric. For a simple non-dispersive onedimensional piezoelectric model of an OHC, the force is related to the displacement and the voltage by 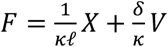, where *κ* is the compliance, *ℓ* is the cell length and *δ* is the piezoelectric strain coefficient. Substituting into Eq. 7 gives the piezoelectric prediction for mechanical power output:

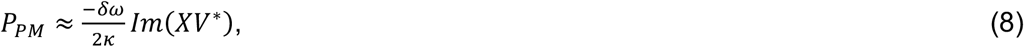

which is identical to Eq. 5 providing 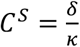 (which is the piezoelectric force coefficient) but derived from the mechanical vs. electrical perspective. Hence, in the simple piezoelectric model electrical power consumed by prestin is equal to the mechanical power driving displacement of the molecular motor and the mechanical load, which satisfies the second law of thermodynamics.

### Mechanical Power Dissipated by the Load

Power consumed by the prestin-motor complex is partially dissipated by heat in the cell itself and partially transmitted to the extrinsic mechanical load -- a load that varies systematically in the cochlea along the tonotopic map. Insight can be gained into OHC power output by my treating the cochlear load as a simple spring-mass-damper. From Newton’s 2nd law: *D^L^X* = *F^T^*, where *X* is displacement of the load and *F^T^* is the total force acting on the load (OHC generated force plus applied force). In the frequency domain, the operator *D^L^* is

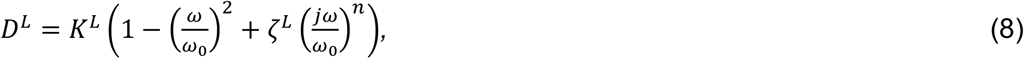

where stiffness of the load and OHC combined is *K^L^*, the passive undamped natural frequency is *ω*_0_ and the nondimensional viscous drag coefficient is *ζ^L^*. Eq. 8 uses a fractional derivative *n* to model the viscous load [27], which gives rise to a power-law frequency dependence of viscous dissipation [10]. For an isolated cell in the dish, *D^L^* describes intrinsic properties of the cell itself plus the media, and in the cochlea *D^L^* describes the addition of the intrinsic properties of the cell plus the cochlear load with frequency, stiffness and damping parameters adjusted accordingly. Multiplying Eq. 8 by velocity and averaging over time gives the power consumed by the viscous load in the frequency domain:

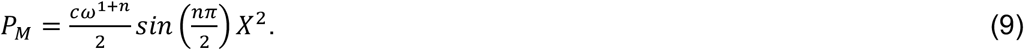

where the damping parameter 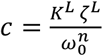. For *n* = 1, which is the standard first approximation, Eq. 9 reduces to the well-known expression for power consumption by a viscous damper: 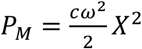. As expected, mass and stiffness do not appear in Eq. 9. Total mechanical power output can easily be estimated from Eq. 9 based on whole-cell displacement in the frequency domain after estimating the viscous dissipation parameters *c* and *n*.

## Results

### Mechanical Power Output Delivered to the Dissipative Load

To determine frequency dependence of prestin motor function in isolated OHCs, mechanical power output was determined from high-frequency voltage-driven displacement data reported by Frank et al. [24] and data reported by Santos-Sacchi & Tan [23]. Fig. 1A illustrates the μ-chamber recording set up, where electromotility was evoked by applying extracellular sinusoidal voltage commands *V_e_* to the base of OHCs. Results for two cells from Frank et al. are shown in Fig. 1 in the form of magnitude (B) and phase (C). Solid curves are fits to the overdamped spring-mass-damper given by Eq. 8 for the longer cell (purple, squares) and the shorter cell (green, circles), with fit parameters: *ω_o_* = (3.45×10^5^, 1.73×10^5^) rad-s^-1^, *ζ^L^* = (1.43,1.74) and *n* = (0.84,0.90) respectively. Mechanical power output was found using Eq. 9, shown as the two solid curves in 1D, peaking at 34 kHz and 73 kHz for the 2 cells. OHC power output was maximum at the frequency when the displacement of the cell lagged the stimulus voltage by ~90°, which is the frequency where the force generated by prestin aligns with cell velocity. Using conventional damping *n* = 1 reduces goodness of the curve fits, but does not change the fact that peak power output occurs at frequencies well above the corner frequency of whole cell displacement. This occurs because the viscous drag force depends on velocity rather than displacement, causing the mechanical power output to reflect velocity squared, in contrast to displacement squared (for *n* = 1).

**Fig. 1.**
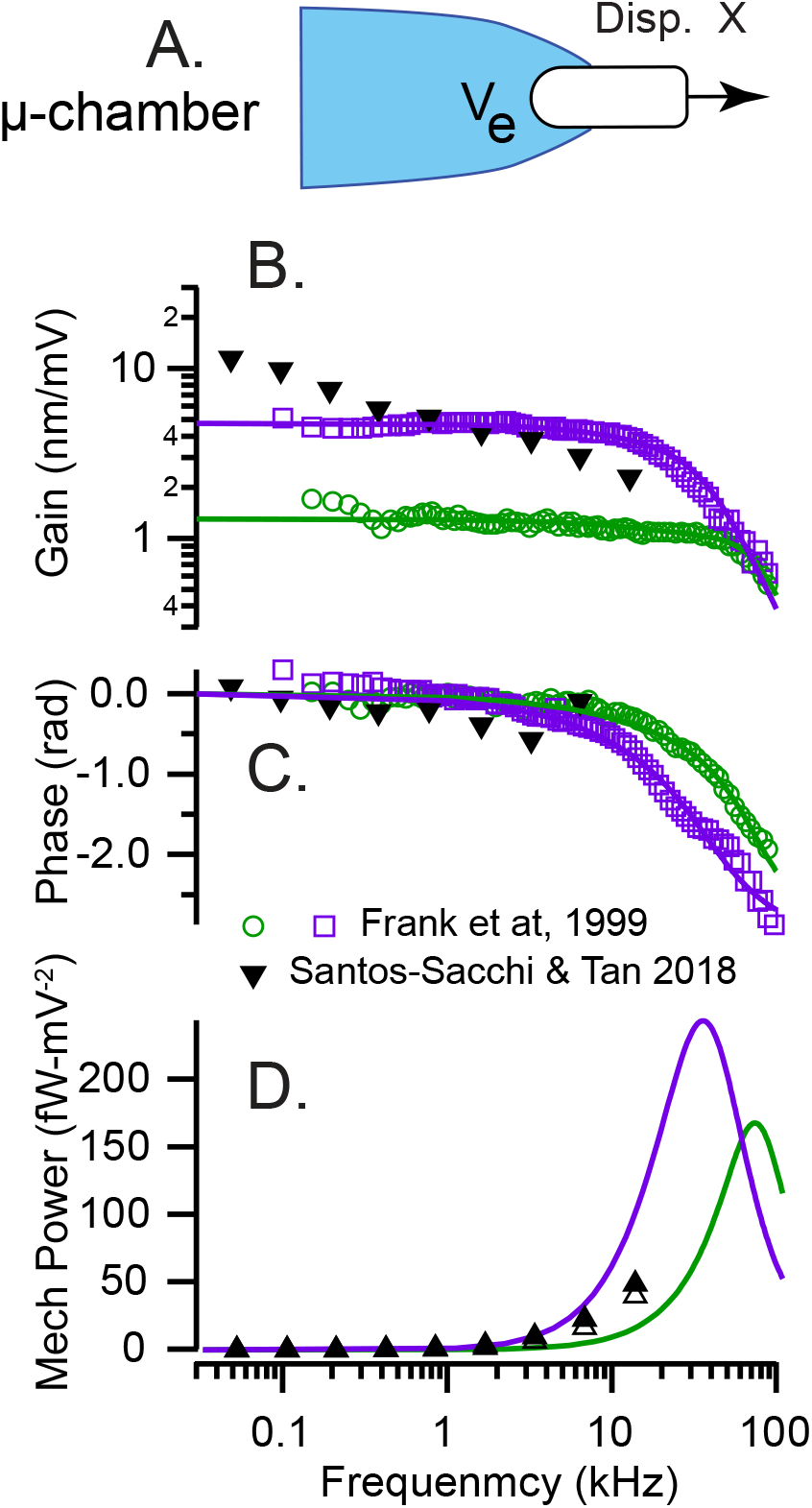
OHC mechanical power output in the dish. A). μ-chamber used to apply sinusoidal voltage stimuli to the base of the cell with the OHC partially extending into media. B-C) Magnitude and phase of the displacement *X* measured by Frank et al. [24] and Santos-Sacchi & Tan [23]. Solid curves are curve-fits to Eq. 8 used to estimate the dissipation parameters “*c*” and “*n*“. D) Mechanical power output was determined from Eq. 9 and measured displacement *X*. Power output computed from the data Frank et al. data peaked at frequencies as high as 74 kHz, when the phase of the displacement was −90° relative to the peak sinusoidal voltage. Power output computed from the Santos-Sacchi & Tan data was continuing to increase up to the highest frequency tested.

For low frequency voltage clamp recordings from OHCs in the dish, voltage is in phase with displacement, and power conversion by the prestin-motor complex is zero (Eq. 5). Hence, low frequency electromotility is only indirectly related to OHC function as a motor. Significant power conversion only occurs at high frequencies as viscosity compels the cell displacement to shift ~90° relative to the voltage, maximizing electrical power consumption *P_E_* and mechanical power output *P_M_*.

In the Frank et al. experiments, the intracellular voltage was not measured, so the precise resting potential *ν_0_* and cycle-by-cycle voltage modulation *ν* were not known. This transmembrane voltage ambiguity likely explains why the isometric force recorded in the μ-chamber configuration (~0.03 nN-mV^-1^ [24]) was smaller than the isometric force recorded in whole-cell voltage clamp (~0.10 nN^-^mV^-1^ [28]). In subsequent μ-chamber experiments by Santos-Sacchi & Tan [23], an offset voltage was applied to restore the resting potential and increase electromotility gain to obtain the summary data reproduced here in Fig. 1B-C (inverted triangles, ▾). Using the Santos-Sacchi & Tan data, Eq. 9 gives the mechanical power output in Fig. 1D. Power was computed directly from their data using the nondimensional OHC viscous drag coefficient estimated for conventional viscous dissipation (*ζ^L^* = 1.43, *n* = 1 open triangles, ▵) and for power law viscous dissipation (*ζ^L^* = 1.43, *n* = 0.84 filled triangles, ▾), with both estimates giving almost identical results. Changing *ζ^L^* relative to the value found from the broader-bandwidth Frank et al. data changes the magnitude of the power output curve but not the frequency dependence. Power output was continuing to increase in the Santos-Sacchi & Tan experiments up to the highest frequency tested (1D, triangles), proving high frequency motor function and forcing rejection of the low-pass hypothesis.

Results in Fig. 1 are consistent with power delivered to a viscous load by a force-driven spring-mass-damper system. Based on simple mechanics, the frequency limit of OHC power output is not limited by the low-pass character of electrically-driven whole-cell displacement, but instead depends strongly on shortening velocity and is limited by the speed at which the prestin-generated force can displace the mechanical load. In the μ-chamber, the mechanical load on the OHC arises from the media and the cell itself. The frequency of maximum real power output is aligned with the best frequency where contributions of mass and stiffness nearly cancel. Ultimately, as computed from Frank et al. data (1D), power output declines at high frequencies as expected from mechanics. The band-pass nature of OHC power output occurs in the μ-chamber because the cell stiffness limits power output at low frequencies, while the fluid and cell mass limit power output at high frequencies. In the cochlea, maximum power output would be expected to occur near the characteristic frequency where the mass and stiffness nearly cancel and the load is dominated by viscous drag [9, 10, 29].

### Electrical Power Converted by the Prestin-Motor ComplexP_P_

To determine the speed of the prestin-motor function in membrane patches, frequency-dependent electrical power consumption was determined directly from macro-patch NLC data reported and by Santos-Sacchi et al. [21], replotted in Fig. 1 as the real *Re*(*C_N_*) (A) and imaginary *Im*(*C^N^*) (B) components. For comparison, capacitance reported by Gale & Ashmore [20] is also shown (right axis, blue circles, real and imaginary components were not separated in the Gale & Ashmore report). The magnitude of the NLC between the two reports differs primarily due to the size of the patch. Both data sets clearly show the magnitude of *C^N^* declining with frequency (2A,B). *Re*(*C^N^*) (2A, thick black) has a low-pass characteristic beginning to roll off below 2kHz and showing a power-law frequency dependence. But, as proven above (Eq. 1, 5), the real component of NLC is not related to electro-mechanical power conversion and therefore has little to do with function of prestin as a motor. Instead, the electrical power consumed by the prestin-motor complex is proportional to negative frequency times the imaginary component of NLC. The Santos-Sacchi et al. data clearly show *Im*(*C^N^*) is indeed negative in prestin-expressing membrane patches and continues to increase in magnitude up to the highest frequency tested (Fig. 2B, thick black).

**Fig. 2.**
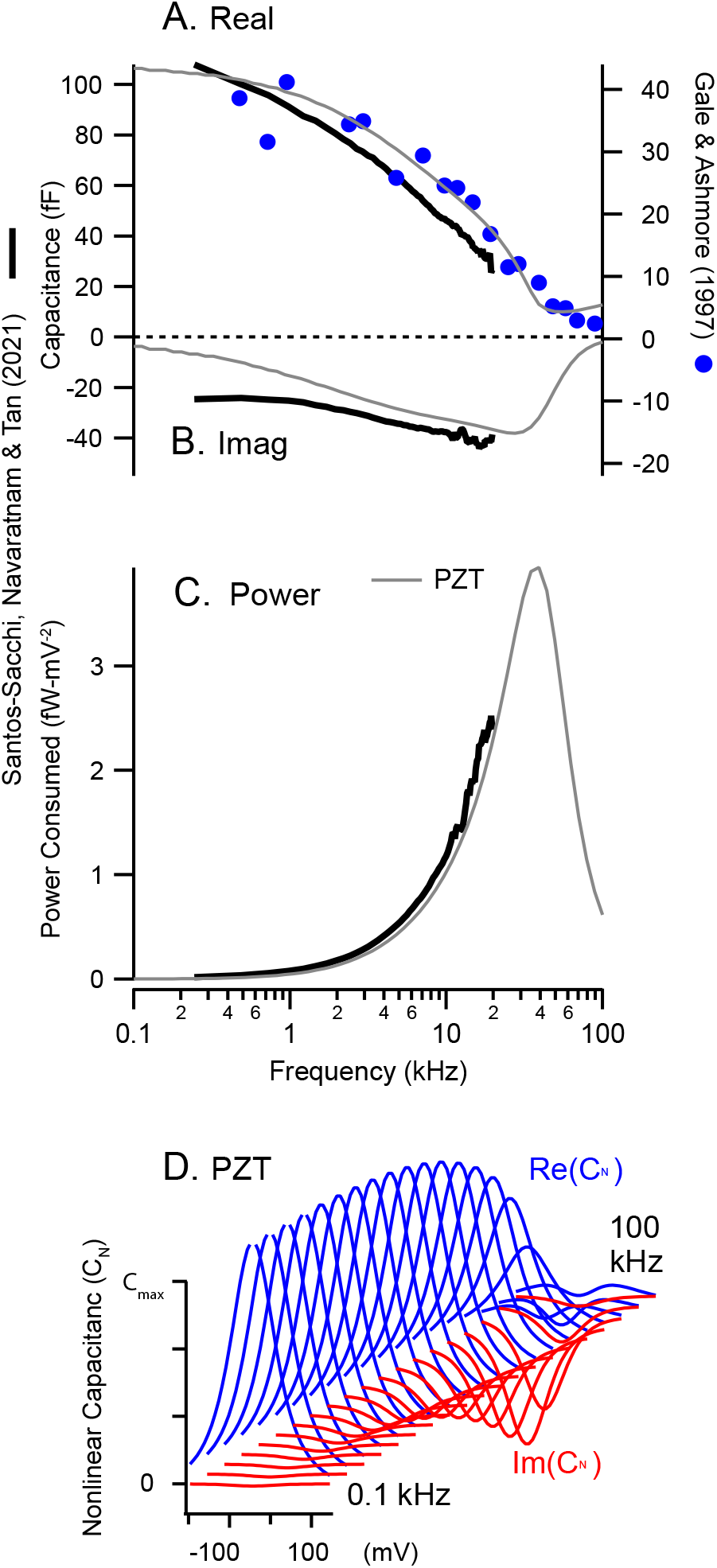
Membrane patch electrical power consumption. A) Real NLC, *Re*(*C^V^*), recorded from OHC membrane macro-patches from data reported by Santos-Sacchi et al. (thick black) [21]. The right axis shows NLC reported by Gale & Ashmore (blue symbols) [20] for comparison. B) Imaginary NLC, *Im*(*C^V^*), recorded from OHC membrane macro-patches from data reported by Santos-Sacchi et al. (thick black). C) Electrical power consumed by the prestin-motor complex computed using Eq. 6 from *Im*(*C^V^*) reported by Santos-Sacchi et al. (thick black). A-C) Thin gray curves are predictions of PZT theory [10]. D) Voltage and frequency dependence of PZT theory corresponding to the solid gray curves in panels A-C.

To interpret electrical power consumed in prestin expressing macropatches, we compared results in Fig. 2 to predictions of OHC piezoelectric theory (PZT [10]). Solid gray curves in Fig. 2A, B and C show theoretical predictions for *Re*(*C^N^*), *Im*(*C^N^*) and *P_P_* respectively. For parameters approximated from the Santos-Sacchi et al. data [21], PZT theory predicts band-pass power consumption to peak near 37 kHz. Results support the hypothesis that electrical power consumed by the prestin motor complex is band-pass and operates at frequencies much higher than would be implied by the low pass character of |*C^W^*|. By the second law of thermodynamics, the electrical power consumed by the membrane patch (2C) must be delivered to dissipation inside the membrane itself and/or to viscosity of the fluid media surrounding the patch (1D).

## Discussion

Results in Fig. 2 demonstrate high-frequency electro-mechanical power conversion by prestin expressing membrane patches, but precisely where the power goes is a subject of debate with hypotheses differing dramatically between reports. In piezoelectric theory applied to isolated OHCs or membrane patches [10] almost all of the power is delivered to the viscous fluid media, and *Im*(*C_n_*) emerges with increasing frequency as viscosity of the fluid viscosity compels the force from prestin to align with the mechanical rate of deformation. Electro-mechanical power conversion in PZT is band-pass, and is predicted to peak under macro-patch voltage clamp conditions near 30 kHz (Fig. 2C, thin gray). The peak occurs at the characteristic frequency of the patch where the load is dominated by viscosity (vs. mass or stiffness). For the same reason, peak power output by OHCs in the cochlea is predicted by PZT to align with characteristic frequency [8–10, 30]. In contrast, if the mechanical load is neglected, *Im*(*C_n_*) arise in transition state theory rate constants, which means the conformation transition itself dissipates all of the power as heat, analogous to dielectric loss [21], with no possibility of power delivery to cochlear amplifier.

Results in Fig. 1 offer compelling evidence that the electrical power consumed by the prestin motor complex is manifested as mechanical power output. In the Frank et al. μ-chamber experiments, mechanical power output was band-pass and peaked between ~30-70 kHz, depending on cell length. Cells continued to output significant power up to the highest frequency tested, 100kHz. Santos-Sacchi et al. improved the μ-chamber approach to control membrane potential, revealing low-pass characteristics of whole-cell displacement, but mechanical power output computed from their data is similar to the Frank et al. report and continued to increase up to the highest frequency tested, 15 kHz. Mechanical power output could not have occurred in these experiments if the imaginary component of NLC recorded membrane patches arises from dielectric loss, supporting the null hypothesis that *Im*(*C_n_*) in voltage clamp experiments (Fig. 2) arises at high frequencies due to the prestin motor complex delivering power to the viscous load.

From the electrical perspective under voltage-clamp conditions, power conversion requires negative *Im*(*C^N^*), which emerges only at high frequencies as *Re*(*C^N^*) becomes small. Although peak *Im*(*C^N^*) is smaller than peak *Re*(*C^N^*) resulting in a frequency dependent decline in the magnitude |*C^W^*|, the decline does not reflect the limiting speed of prestin motor function or a decline in electro-mechanical power conversion. Emergence of *Im*(*C^N^*) under voltage clamp reveals a shift in the phase of the prestin-dependent charge displacement relative to force, which is in phase with *dν*/*dt* at low frequencies (giving rise to real NLC) and in phase with *ν* at high frequencies (giving rise to negative imaginary NLC). The imaginary NLC appears in electrophysiological recordings as a real-valued admittance [31], but it is a reversible charge displacement that delivers power to the mechanical load in contrast to a conduction current or dielectric loss that dissipates power as heat. From the mechanical perspective, emergence of *Im*(*C^N^*) under voltage clamp conditions reflects alignment of the force with shortening velocity, which is requisite for OHCs in the cochlea to deliver mechanical power to the viscous cochlear load.

In summary, present results are consistent with the hypothesis that the speed of force generation by the prestin-motor complex is nearly instantaneous, while the speed of conformational change is limited by the load against which the protein must deform. For maximum power output, the timing of prestin-motor charge displacement during the power stroke must be shifted −90° relative to the charge displacement measured at low frequencies in the absence of significant mechanical load. The phase shift is reflected on both the electrical and mechanical sides of the motor. Mechanical power output to a dissipative viscous load requires force to be in phase with velocity. The present report only analyzed data collected under voltage-clamp commands. Nevertheless, the above analysis shows electro-mechanical power conversion is maximized only when voltage, current, force and velocity occur at specific phases relative to each other -- requirements likely met by setting the mechano-electrical transduction current, RC corner frequency, cell stiffness, and level of prestin expression along the tonotopic map in the cochlea [29, 32–34]. Electrical properties of the Corti are also implicated as important in power conversion, through the influence of the electro-anatomy on frequency-dependent extracellular potentials and OHC transmembrane voltage [35].

## Acknowledgement

Support was provided by the National Institutes on Deafness and other Communication Disorders grant R01 DC006685.

## Citations

[1] Brownell, W.E., Bader, C.R., Bertrand, D. & de Ribaupierre, Y. 1985 Evoked mechanical responses of isolated cochlear outer hair cells. Science 227, 194–196. (doi:10.1126/science.3966153).

[2] Liberman, M.C., Gao, J., He, D.Z., Wu, X., Jia, S. & Zuo, J. 2002 Prestin is required for electromotility of the outer hair cell and for the cochlear amplifier. Nature 419, 300–304. (doi:10.1038/nature01059).

[3] Zheng, J., Shen, W., He, D.Z., Long, K.B., Madison, L.D. & Dallos, P. 2000 Prestin is the motor protein of cochlear outer hair cells. Nature 405, 149–155. (doi:10.1038/35012009).

[4] Dong, X.X., Ospeck, M. & Iwasa, K.H. 2002 Piezoelectric reciprocal relationship of the membrane motor in the cochlear outer hair cell. Biophys J 82, 1254–1259. (doi:10.1016/S0006-3495(02)75481-7).

[5] Yang, J.S. & Batra, R.C. 1994 A Theory of Electroded Thin Thermopiezoelectric Plates Subject to Large Driving Voltages. J Appl Phys 76, 5411–5417. (doi:Doi 10.1063/1.358489).

[6] Tolomeo, J.A. & Steele, C.R. 1995 Orthotropic piezoelectric properties of the cochlear outer hair cell wall. J Acoust Soc Am 97, 3006–3011. (doi:10.1121/1.411865).

[7] Mountain, D.C. & Hubbard, A.E. 1994 A piezoelectric model of outer hair cell function. J Acoust Soc Am 95, 350–354. (doi:10.1121/1.408273).

[8] Meaud, J. & Grosh, K. 2012 Response to a pure tone in a nonlinear mechanical-electrical-acoustical model of the cochlea. Biophys J 102, 1237–1246. (doi:10.1016/j.bpj.2012.02.026).

[9] O Maoiléidigh, D. & Hudspeth, A.J. 2013 Effects of cochlear loading on the motility of active outer hair cells. Proc Natl Acad Sci U S A 110, 5474–5479. (doi:10.1073/pnas.1302911110).

[10] Rabbitt, R.D. 2020 The cochlear outer hair cell speed paradox. Proc Natl Acad Sci U S A. (doi:10.1073/pnas.2003838117).

[11] Ge, J., Elferich, J., Dehghani-Ghahnaviyeh, S., Zhao, Z., Meadows, M., von Gersdorff, H., Tajkhorshid, E. & Gouaux, E. 2021 Molecular mechanism of prestin electromotive signal amplification. Cell 184, 4669–4679 e4613. (doi:10.1016/j.cell.2021.07.034).

[12] Bavi, N., Clark, M.D., Contreras, G.F., Shen, R., Reddy, B.G., Milewski, W. & Perozo, E. 2021 Prestin’s conformational cycle underlies outer hair cell electromotility. Nature.(doi:10.1038/s41586-021-04152-4).

[13] Butan, C., Song, Q., Bai, J.P., Tan, W.J.T., Navaratnam, D. & Santos-Sacchi, J. 2022 Single particle cryo-EM structure of the outer hair cell motor protein prestin. Nat Commun 13, 290. (doi:10.1038/s41467-021-27915-z).

[14] Dewey, J.B., Altoe, A., Shera, C.A., Applegate, B.E. & Oghalai, J.S. 2021 Cochlear outer hair cell electromotility enhances organ of Corti motion on a cycle-by-cycle basis at high frequencies in vivo. Proc Natl Acad Sci U S A 118. (doi:10.1073/pnas.2025206118).

[15] Cooper, N.P., Vavakou, A. & van der Heijden, M. 2018 Vibration hotspots reveal longitudinal funneling of sound-evoked motion in the mammalian cochlea. Nat Commun 9, 3054. (doi:10.1038/s41467-018-05483-z).

[16] Dewey, J.B., Applegate, B.E. & Oghalai, J.S. 2019 Amplification and Suppression of Traveling Waves along the Mouse Organ of Corti: Evidence for Spatial Variation in the Longitudinal Coupling of Outer Hair Cell-Generated Forces. J Neurosci 39, 1805–1816. (doi:10.1523/JNEUROSCI.2608-18.2019).

[17] Wang, Y., Steele, C.R. & Puria, S. 2016 Cochlear Outer-Hair-Cell Power Generation and Viscous Fluid Loss. Sci Rep 6, 19475. (doi:10.1038/srep19475).

[18] Reuter, G., Kossl, M., Hemmert, W., Preyer, S., Zimmermann, U. & Zenner, H.P. 1994 Electromotility of outer hair cells from the cochlea of the echolocating bat, Carollia perspicillata. J Comp Physiol A 175, 449–455. (doi:10.1007/BF00199252).

[19] Vavakou, A., Cooper, N.P. & van der Heijden, M. 2019 The frequency limit of outer hair cell motility measured in vivo. Elife 8. (doi:10.7554/eLife.47667).

[20] Gale, J.E. & Ashmore, J.F. 1997 The outer hair cell motor in membrane patches. Pflugers Arch 434, 267–271. (doi:10.1007/s004240050395).

[21] Santos-Sacchi, J., Navaratnam, D. & Tan, W.J.T. 2021 State dependent effects on the frequency response of prestin’s real and imaginary components of nonlinear capacitance. Sci Rep 11, 16149. (doi:10.1038/s41598-021-95121-4).

[22] Santos-Sacchi, J. & Tan, W. 2019 Voltage Does Not Drive Prestin (SLC26a5) Electro-Mechanical Activity at High Frequencies Where Cochlear Amplification Is Best. iScience 22, 392–399. (doi:10.1016/j.isci.2019.11.036).

[23] Santos-Sacchi, J. & Tan, W. 2018 The Frequency Response of Outer Hair Cell Voltage-Dependent Motility Is Limited by Kinetics of Prestin. J Neurosci 38, 5495–5506. (doi:10.1523/JNEUROSCI.0425-18.2018).

[24] Frank, G., Hemmert, W. & Gummer, A.W. 1999 Limiting dynamics of high-frequency electromechanical transduction of outer hair cells. Proc Natl Acad Sci U S A 96, 4420–4425. (doi:10.1073/pnas.96.8.4420).

[25] Sasmal, A. & Grosh, K. 2019 Unified cochlear model for low- and high-frequency mammalian hearing. Proc Natl Acad Sci U S A 116, 13983–13988. (doi:10.1073/pnas.1900695116).

[26] Mosgaard, L.D., Zecchi, K.A. & Heimburg, T. 2015 Mechano-capacitive properties of polarized membranes. Soft Matter 11, 7899–7910. (doi:10.1039/c5sm01519g).

[27] Freed, A.D. & Diethelm, K. 2006 Fractional calculus in biomechanics: a 3D viscoelastic model using regularized fractional derivative kernels with application to the human calcaneal fat pad. Biomech Model Mechanobiol 5, 203–215. (doi:10.1007/s10237-005-0011-0).

[28] Iwasa, K.H. & Adachi, M. 1997 Force generation in the outer hair cell of the cochlea. Biophys J 73, 546–555. (doi:10.1016/S0006-3495(97)78092-5).

[29] Bidone, T. & Rabbitt, R. 2022 The perfect match of outer hair cells in the cochlea. In review.

[30] Nam, J.H. & Fettiplace, R. 2010 Force transmission in the organ of Corti micromachine. Biophys J 98, 2813–2821. (doi:10.1016/j.bpj.2010.03.052).

[31] Farrell, B., Ugrinov, R. & Brownell, W. 2006 Frequency dependence of admittance and conductance of the outer hair cell. In Auditory Mechanisms, Processes and Models (eds. A. Nuttall, T. Ren, K. Gillespie, K. Grosh & E. de Boer), pp. 231–232. New Jersey, World Scientific.

[32] Nam, J.H. & Fettiplace, R. 2012 Optimal electrical properties of outer hair cells ensure cochlear amplification. PLoS One 7, e50572. (doi:10.1371/journal.pone.0050572).

[33] Altoe, A. & Shera, C.A. 2022 The outer-hair-cell RC time constant: A feature, not a bug, of the mammalian cochlea. bioRxiv. (doi:https://doi.org/10.1101/2022.02.02.478769).

[34] Lukashkin, A.N. & Russell, I.J. 1997 The voltage dependence of the mechanoelectrical transducer modifies low frequency outer hair cell electromotility in vitro. Hear Res 113, 133–139. (doi:10.1016/s0378-5955(97)00135-4).

[35] Lukashkina, V.A., Levic, S., Lukashkin, A.N., Strenzke, N. & Russell, I.J. 2017 A connexin30 mutation rescues hearing and reveals roles for gap junctions in cochlear amplification and micromechanics. Nat Commun 8, 14530. (doi:10.1038/ncomms14530).

